# Excitatory synaptic transmission is differentially modulated by opioid receptors along the claustro-cingulate pathway

**DOI:** 10.1101/2025.03.31.646444

**Authors:** Jacob M. Reeves, Erwin Arias-Hervert, Gracianne E. Kmiec, William T. Birdsong

## Abstract

The anterior cingulate cortex (ACC) plays a pivotal role in processing pain and emotion, communicating with both cortical and subcortical regions involved in these functions. The claustrum (CLA), a subcortical region with extensive connectivity to the ACC also plays a critical role in pain perception and consciousness. Both ACC and CLA express Kappa (KOR), Mu (MOR), and Delta (DOR) opioid receptors, yet whether and how opioid receptors modulate this circuit is poorly understood. This study investigates the effects of opioid receptor activation on glutamatergic signaling in CLA-ACC circuitry using spatial transcriptomics, slice electrophysiology, optogenetics, and pharmacological approaches in mice. Our results demonstrated that excitatory inputs generated by the CLA onto layer 5 pyramidal cells (L5 PYR) in the ACC are reduced by KOR, MOR, and DOR agonists. However, only KOR agonists reduce monosynaptic transmission from the CLA onto L5 ACC PYR cells, highlighting the unique role of KOR in modulating the CLA-ACC pathway. MOR agonists had a heterogeneous effect on optically-evoked excitatory postsynaptic currents (oEPSCs), significantly reducing longer-latency excitatory responses while only modestly inhibiting the short latency excitatory postsynaptic currents. DOR agonists only reduce slower, longer-latency recurrent excitatory responses. These findings provide new insights into how opioid receptors regulate the claustro-cingulate circuit and demonstrate the distinct, receptor-specific modulation of synaptic transmission within this network.

## Introduction

The ACC is a critical hub for emotional and pain-related processes (Barthas et al., 2015). As such, the ACC both sends and receives information from cortical and subcortical regions involved in pain perception and emotional regulation (Krieghoff et al., 2011, Rolls 2019, Shi et al., 2022). Within these pathways, opioid peptides can activate KOR, MOR, and DOR opioid receptors to decrease neuronal activity and neurotransmission and ultimately modulate circuit function. Local ACC neurons express MOR and DOR while also receiving MOR-sensitive innervation from thalamic inputs (Mansour et al., 1987, Mansour et al., 1994, Vogt et al., 1995, Birdsong et al., 2019, Zamfir et al., 2023). While it has been shown that the prefrontal cortex (PFC) receives KOR-sensitive inputs from the basolateral amygdala (Yarur et al., 2024) and the ventral tegmental area (Margolis et al., 2006), no KOR-sensitive inputs into the ACC or KOR-dependent modulation of local excitatory processing have been characterized.

The CLA is a subcortical brain region lying between the insula and the lateral striatum that expresses high levels of KOR, moderate levels of MOR, and low levels of DOR (Tempel and Zukin 1987, George et al., 1994, Crick and Koch 2005, Wang et al., 2011, Chen et al., 2020). The CLA exhibits extensive cortical connectivity; the region’s activity is associated with processes including pain perception, attention, consciousness, memory, and sleep (Atlan et al., 2024, Smith et al., 2020). The CLA sends dense projections to the ACC and other frontal cortical areas (Jackson et al., 2018, Ährlund-Richter et al., 2019). While a role for the CLA-ACC pathway in pain processing has been highlighted (Xu et al., 2022, Ntamati et al., 2023, Zhang and Zamponi 2024), whether and how opioids modulate the CLA-ACC pathway is understudied. Several studies have highlighted opioid action in CLA-prefrontal circuits. Chronic social defeat stress attenuated the excitatory output of the CLA-prelimbic cortex circuit through dynorphin/KOR signaling, being critical for depressive-related behaviors in male mice (Wang et al., 2023). Salvinorin A, a KOR agonist, increased functional connectivity between the CLA and ACC in humans suggesting that KOR agonists can modulate CLA-ACC circuitry (Bagdasarian et al., 2024). However, direct physiological studies investigating (1) the effects of KOR activation on CLA-ACC networks, (2) the location of KOR within these networks, and (3) whether MOR and DOR activation also modulates these networks are lacking and are the focus of this study.

Here we employed spatial transcriptomics, slice electrophysiology, optogenetics, and pharmacology to characterize opioid receptor modulation along the CLA-ACC pathway in mice. We found that stimulation of CLA inputs in the ACC elicited excitatory transmission with varying latencies and complex waveforms, suggesting CLA inputs elicit both direct and recurrent excitation in the ACC. KOR, MOR, and DOR agonists all reduced excitatory transmission but to varying degrees. When recurrent polysynaptic transmission was blocked, only KOR agonists retained their ability to inhibit CLA-ACC excitatory transmission. Therefore, monosynaptic inputs from the CLA were regulated by KOR, while MOR and DOR inhibition was restricted to recurrent cortical excitatory circuits. Together, our data reveal unique roles for KOR, MOR, and DOR in regulating the CLA-ACC pathway. These data illustrate how the claustro-cingulate circuit is modulated by opioids and how different opioid receptor subtypes can independently modulate neuronal communication at the circuit level.

## Methods

Animals. All procedures were conducted in accordance with the National Institutes of Health guidelines and with approval from the Institutional Animal Care and Use Committee at the University of Michigan. Mice were maintained on a 12 hr light/dark cycle and given ad libitum access to food and water. C57Bl/6J mice were obtained from Jackson Laboratories. Mice were postnatal day 40-80 at the time of viral injection and postnatal day 55-110 at the time of brain slice preparation. Mice of both sexes were used.

Spatial Transcriptomics. Publicly available data from the Allen Institute’s - “Allen Brain Cell Atlas” was used to identify opioid receptor transcripts in the CLA (Allen Brain Cell Atlas (RRID:SCR_024440) https://portal.brain-map.org/atlases-and-data/bkp/abc-atlas, Yao et al., 2023). The MERSIFH-C57BL6J-638850 with Imputed Genes + Reconstructed Coordinates dataset was used. Brain Slice IDs C57BL6J-638850.52 and C57BL6J-638850.54 were used to obtain expression data results from the CLA, bilaterally. The neurotransmitter type, Glut, was selected to isolate glutamate expressing cells in the CLA and oprk1, oprm1, and oprd1 genes were selected. Because gene expression data are reported in binned ranges, all cells from each expression range were randomly assigned a discrete expression value that fell within their specific assigned range using a random number generator in Microsoft Excel. Expression values were then plotted on a violin plot as Log2(CPM+1).

Stereotaxic injections. Stereotaxic injections were performed as previously described (Jaeckel et al., 2024) to deliver adeno-associated virus (AAV) to express channelrhodopsin (ChR2) (Gift of Karl Deisseroth produced by UNC Vector Core). Mice were injected bilaterally with an adeno-associated virus type 2 encoding channelrhodopsin-2 [ChR2; AAV2-hsyn-ChR2(H134R)-EYFP] targeting the claustrum (CLA). In total, ∼ 100 nL of virus was injected into the CLA (A/P, + 1.5 mm; M/L, ± 2.7 mm; D/V, 3.4-3.6 mm). To target CLA projecting ACC neurons, mice were injected bilaterally with a retro cre-dependent adeno-associated virus (AAVrg-EF1a-Cre) in the ACC (A/P, + 0.75 mm; M/L, ± 0.4 mm; D/V, 1.55-1.65 mm) and (AAV5-EF1a-double floxed-hChR2(H134R)-EYFP-WPRE-HGHpA) in the CLA. ∼ 100 nL was injected into both regions.

Brain slice electrophysiology. Brain slices were prepared 2–3 weeks following injection of ChR2. Mice were deeply anesthetized with isoflurane and decapitated. Brains were removed and mounted for slicing with a vibratome (Model 7000 smz, Campden Instruments). During slicing, brains were maintained at 34°C in carbogenated Krebs’ solution containing the following (in mM): 136 NaCl, 2.5 KCl, 1.2 MgCl2–6H2O, 2.4 CaCl2–2 H2O, 1.2 NaH2PO4, 21.4 NaHCO3, 11.1 dextrose supplemented with 5 µM MK-801. 300 µM coronal sections containing the ACC were made and incubated in carbogenated Krebs’ solution supplemented with 10 µM MK-801 at 32°C for 30 min. Slices were then maintained at room temperature in carbogenated Krebs’ solution until used for recording. Only one cell was recorded from each slice due to pharmacological drug washes. For ACC recordings, borosilicate glass patch pipettes (Sutter Instrument) were pulled to a resistance of 2.0–3.0 MΩ and filled with a Cesium gluconate-low chloride based internal solution (in mM: 135 Cesium Gluconate, 1 EGTA, 1.5 MgCl2, 10 HEPES (Na), 2 Na ATP, 0.4 Mg GTP, 7.8 Na2 phosphocreatine). Slices were placed in the recording chamber and continuously perfused with carbogenated Krebs’ solution at 32–33°C. Layer 5 pyramidal neurons were identified based on cell morphology. Whole-cell voltage clamp recordings were made in layer 5 pyramidal neurons at −65 mV holding potential to isolate excitatory postsynaptic currents (EPSCs). Cells were dialyzed for 5 mins before experimental recordings. oEPSCs were elicited every 30 s by LED illumination through the microscope objective (Olympus BX51WI) using a transistor-transistor logic (TTL)-controlled LED driver and a 470 nm LED (Thorlabs). The LED stimulation duration was 1 ms and power output measured after the microscope objective ranged from 0.83 to 5.59 mW, adjusted to obtain consistent current amplitudes across cells. For TTX + 4-AP recordings (Fig. 3), baseline oEPSC values smaller than 150 pA were discarded. oEPSCs were recorded before moving to the next drug condition (baseline, agonist, and antagonist). Whole-cell recordings were made with a MultiClamp 700B amplifier (Molecular Devices) digitized at 20 kHz (National Instruments BNC-2090A). Synaptic recordings were acquired using Matlab WaveSurfer (MathWorks). Series resistance was monitored throughout the recordings, and only recordings in which the series resistance remained <10 MΩ and did not increase more than 20% were included.

Analysis of Electrophysiology Data. Raw data were analyzed using Axograph. For each condition (baseline, agonist, and antagonist), baseline subtracted sweeps were filtered with a 1 kHz low-pass gaussian filter and averaged together and plotted. For the agonist and antagonist drug conditions, the first 4 sweeps were omitted from the average to allow for equilibration of drug and washout of drug within the tissue. Peak current amplitude was calculated from the maximum value of the averaged traces and plotted as % reduction. Cells that have no connecting line for the antagonist (Fig. 1, 2) either died or had a series resistance too high to include in this study during the antagonist wash. Charge transfer was analyzed by calculating the area of the oEPSC at time point 0 - 50 ms in relation to light flash and was plotted as % reduction. For event detection analysis, the 1st derivative was taken of each sweep, and an event amplitude threshold analysis was performed (settings: minimum event separation = 0 (ms), capture baseline = 1 (ms), capture length = 5 (ms), time to peak for typical event = 0.8 (ms) (EPSC), peak measurement interval = 0 (ms)). The latency was calculated from the time of the light flash and events were detected from -50 - 50 ms in relation to the light flash.

**Figure 1:**
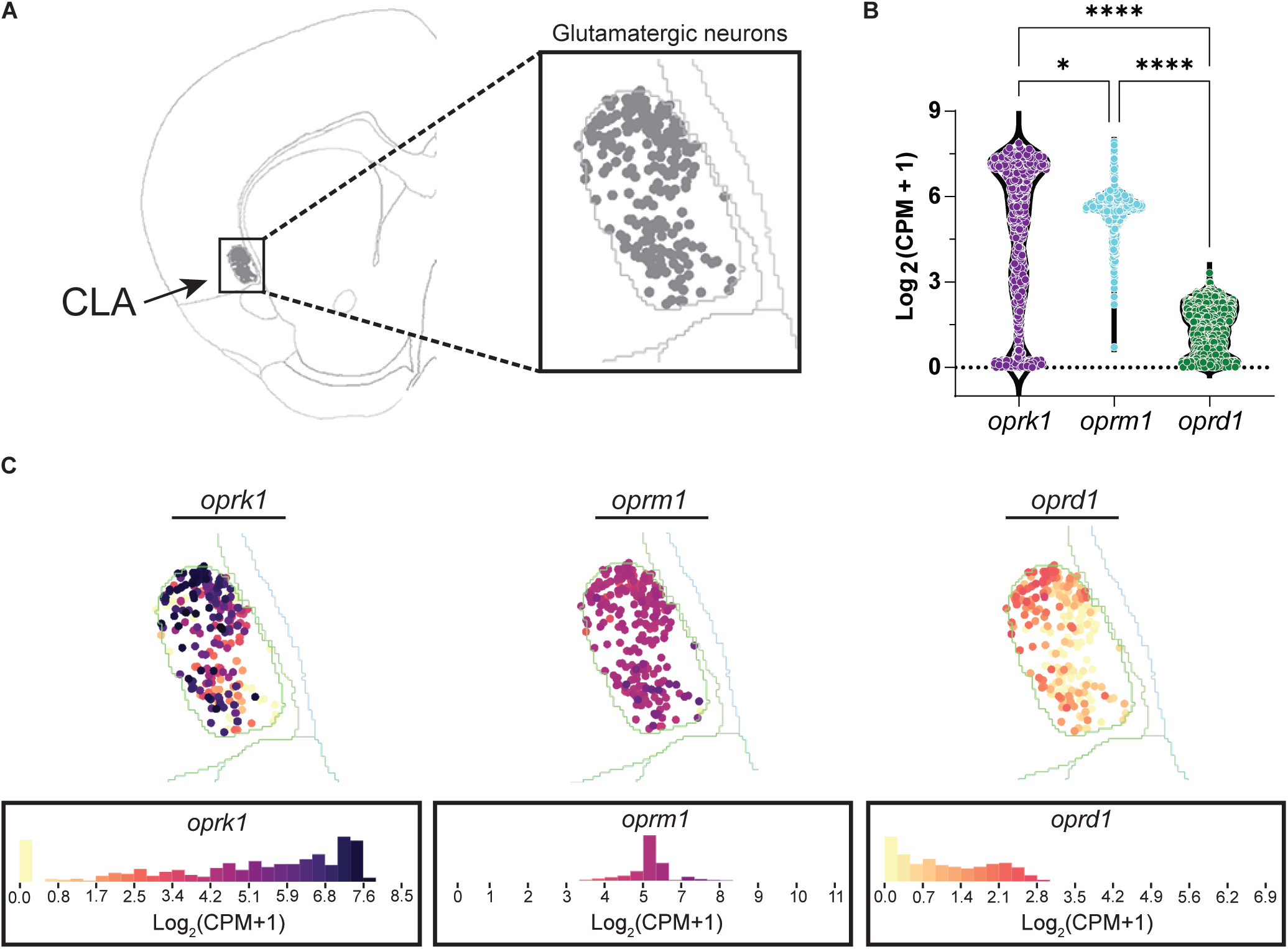
Kappa, Mu, and Delta transcripts are differentially expressed in CLA glutamatergic neurons. A, Example image of a coronal slice of a mouse brain from the Allen Institute’s ABC Atlas - Whole mouse brain (Dataset: MERSIFH-C57BL6J-638850 with Imputed Genes + Reconstructed Coordinates; Brain Slice ID: C57BL6J-638850.52). Grey individual dots are glutamate expressing neurons highlighted in the CLA. B, Violin plot of summary data of oprk1, oprm1, and oprd1 transcripts in CLA glutamate expressing neurons, oprk1: cells = 1020; oprm1: cells = 1026; oprd1: cells = 1025; N = 1, n (slices) = 2; oprk1 vs. oprm1, p = 0.0311; oprk1 vs. oprd1, p = <0.0001; oprm1 vs. oprd1, p = <0.0001; Kruskal-Wallis test with Dunn’s multiple comparisons test). Each dot represents an individual cell. C, Example images of oprk1, oprm1, and oprd1 expression in glutamate expressing cells in the CLA. A histogram heatmap of the expression profiles for each transcript is below the example image.

**Figure 2:**
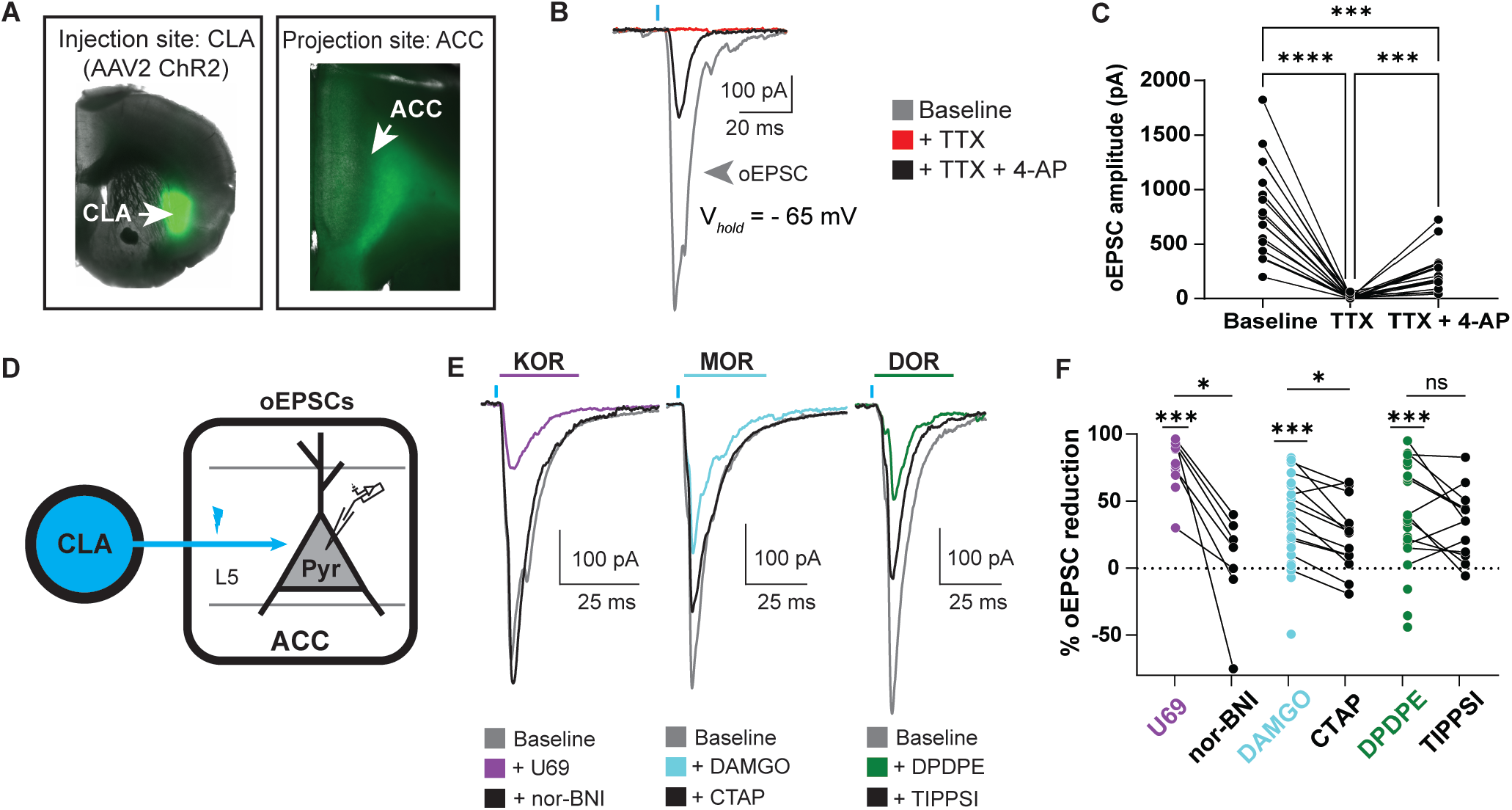
KOR, MOR, and DOR suppress CLA-evoked excitatory synaptic transmission onto L5 ACC PYR cells. A, Channelrhodopsin was virally transduced into the CLA of WT mice; ChR2 expression is seen in CLA terminals innervating the ACC. B, Example oEPSC recorded under baseline conditions (grey) in TTX (1 µM) (red), and in TTX (1 µM) + 4AP (black). C, Summary data of oEPSCs of all recordings shown in (B) with responses plotted at raw oEPSC amplitude in picoamps (pA) (Baseline vs. TTX, p = <0.0001; Baseline vs. TTX + 4-AP, p = 0.0001; TTX vs. TTX +4-AP, p = 0.0002; Repeated measures one-way ANOVA with Šídák’s multiple comparisons tests; N = 9, n = 18). D, Schematic of recording oEPSCs of L5 ACC PYR cells evoked by optical stimulation of CLA input. E, Example traces of oEPSCs elicited from optical stimulation of the CLA terminals in the ACC during baseline (gray), application of U69 (1 µM, left panel, purple), followed by nor-BNI (1 µM, left panel, black), or application of DAMGO (1 µM, middle panel, cyan) followed by CTAP (1 µM, middle panel, black), or DPDPE (1 µM, right panel, green) followed by TIPPSI (1 µM, right panel, black). Blue bars: 1 ms of 470 nm light stimulation. F, Summary data of oEPSCs of all recordings as shown in (E) with responses plotted as a percent oEPSC reduction relative to baseline oEPSC amplitudes: Baseline vs. U69, p = 0.0003, N = 11, n = 11; U69 vs. norBNI, p = 0.0205, N = 11, n = 7; Baseline vs. DAMGO, p = 0.0002, N = 20, n = 26; DAMGO vs. CTAP, p = 0.0032, N = 20, n = 14; Baseline vs. DPDPE, p = 0.0004, N = 17, n = 25; DPDPE vs. TIPPSI, p = 0.2562, N = 17, n = 14. Mixed-effects analysis with Šídák’s multiple comparisons tests conducted with raw amplitudes. *p ≤ 0.05; ** p ≤ 0.01; *** p ≤ 0.001; **** p < 0.0001. Data are described as the mean ± standard error of the mean.

Statistics. Statistical analyses were performed using GraphPad Prism (GraphPadSoftware Inc., San Diego, CA). Statistical comparisons were made using a t-test (ratio or unpaired) or one-way or two-way ANOVA with Šídák’s multiple comparisons tests or Kruskal-Wallis test with Dunn’s multiple comparisons test. Detailed statistical descriptions are reported in each respective figure legend. For all experiments, statistical significance was defined as p < 0.05. For all comparisons, n (number of cells) and N (number of animals) are both reported.

## Results

### Oprk1 exhibits dense expression, oprm1 exhibits moderate expression, and oprd1 exhibits low expression in CLA glutamatergic neurons

Previous studies have highlighted that the KOR mRNA transcript, oprk1, is highly expressed in the rat CLA while the MOR mRNA transcript, oprm1, exhibits moderate expression, and the DOR mRNA transcript, oprd1, exhibits low expression (George et al., 1994, Mansour et al., 1994). However, the specific cell types that express these opioid transcripts are not identified in those studies. In this study, we wanted to observe the expression profiles of oprk1, oprm1, and oprd1 in glutamatergic neurons of the CLA. To answer this question, we used publicly available multiplexed error-robust fluorescence in situ hybridization (MERFISH) data from the Allen Institute to identify glutamate expressing neurons in the CLA and observe the expression values of each opioid receptor transcript (Allen Brain Cell Atlas (RRID:SCR_024440)). Using the dataset (MERSIFH-C57BL6J-638850 with Imputed Genes + Reconstructed Coordinates, Kunst et al., 2023), we isolated cells in the CLA that were transcriptionally classified as glutamatergic (Fig. 1A). We identified and evaluated coronal brain slices from this dataset that aligned with our CLA stereotactic injection coordinates (brain Slice IDs: C57BL6J-638850.52 and C57BL6J-638850.54). Across these glutamatergic CLA neurons, oprk1 was found to be the most densely expressed opioid receptor transcript, oprm1 exhibited consistent moderate expression, while oprd1 expression was the lowest (in Log2(CPM+1): 25% percentile, oprk1: 3.1, oprm1: 5.3, and oprd1: 0.6; 75% percentile, oprk1: 7.1, oprm1: 5.8, and oprd1: 3.3; Fig 1B, C). These results suggest that CLA glutamate neurons express high, dense levels of KOR mRNA transcript, moderate levels of MOR mRNA transcript, and low levels of DOR mRNA transcript.

### Opioid receptor agonists suppress excitatory synaptic transmission generated by CLA inputs onto L5 ACC Pyramidal cells

Previous studies have established that glutamatergic CLA neurons innervate the ACC, with both regions showing dense opioid receptor expression (Tempel and Zukin 1987, George et al., 1994, Chia et al., 2017, 2020, Ntamati et al., 2023); however, little is known about the opioid sensitivity of the CLA projections to the ACC. Because of the dense KOR expression, moderate MOR expression, and low DOR expression within the CLA, we hypothesized that excitatory synaptic transmission from CLA to ACC would be reduced by KOR and MOR agonists but not DOR agonists. We tested the opioid sensitivity of ACC-projecting CLA neurons by injecting the blue light-sensitive channelrhodopsin (AAV2-ChR2 (H134R)) into the CLA of wildtype (WT) mice (Fig. 2A). Whole-cell patch-clamp recordings were made from L5 ACC PYR cells in acute mouse brain slice preparations. Optical stimulation of CLA terminals in the ACC generated an oEPSC that was blocked by TTX and exhibited variable recovery with TTX + 4-AP (oEPSC amplitude (picoamps = pA); Baseline: 832.5 ± 98.8 pA; TTX: 14.02 ± 3.9 pA; TTX + 4-AP: 242.0 ± 43.0 pA, Fig. 2B, C). Together these data confirm that the L5 ACC PYR cells receive excitatory synaptic transmission from the CLA.

Prior studies have shown that the ACC is innervated by the CLA but not by other brain regions surrounding the CLA (Qadir et al., 2018, Atlan et al., 2024). To further verify that our viral injection paradigm results in optical activation of predominantly CLA inputs to the ACC, we injected a retrograde cre-expressing virus (AAVrg-EF1a-Cre) in the ACC and a cre-dependent channelrhodopsin (AAV5-double-floxed-hChR2-EYFP) into the CLA and surrounding areas. Fluorescence expression was only localized to the CLA and not surrounding areas, and CLA axon terminals labeled with YFP were abundant in the ACC (Fig. S1). These data suggest that oEPSCs recorded in the ACC following injection of AAV2-ChR2 into the CLA are predominantly from excitation of CLA inputs rather than the result of viral leakage into surrounding areas that project to the ACC.

Next, we investigated the opioid sensitivity of excitatory responses elicited by optical stimulation of CLA inputs in the ACC. The KOR-selective agonist U69,593 (U69) reduced the oEPSC amplitude, and the KOR-selective antagonist nor-BNI largely recovered the oEPSC (% oEPSC reduction; U69: 80.3 ± 4.1%, nor-BNI: 3.5 ± 14.6%, Fig. 2E, F). The MOR-selective agonist DAMGO reduced the oEPSC amplitude, and the MOR-selective antagonist CTAP recovered the oEPSC amplitude (% oEPSC reduction; DAMGO: 32.2 ± 6.1%, CTAP: 11.4 ± 13.9%, Fig. 2E, F). The DOR-selective agonist DPDPE reduced the oEPSC amplitude, while the DOR-selective antagonist TIPPSI did not recover the oEPSC amplitude (% oEPSC reduction; DPDPE: 39.0 ± 7.8%, TIPPSI: 33.7 ± 7.0%, Fig. 2E, F). There was no effect on sex across drug conditions, so we combined sexes for this study (oEPSC; U69: p = 0.2051, DAMGO: p = 0.7059, DPDPE: p = 0.8604; two-way ANOVA). These data suggest that KOR, MOR, and DOR are functionally present in the CLA-ACC circuit and reduce excitatory synaptic transmission.

**Supplementary Figure 1.**
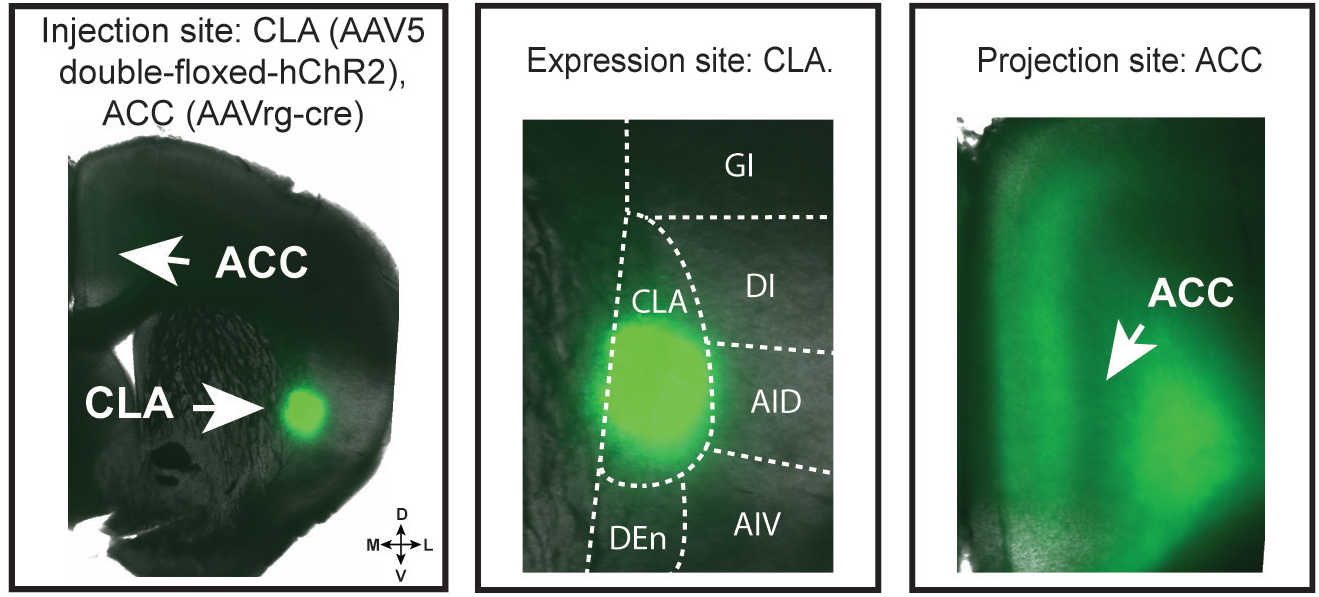
The ACC is innervated by CLA and not surrounding brain regions. Cre-dependent ChR2 (AAV5-EF1a-double floxed-hChR2(H134R)-EYFP-WPRE-HGHpA) was injected in the CLA and retrograde cre-expressing virus (AAVrg-cre) was injected in the ACC. The middle and right images are higher magnification views of the image on the left. Expression is localized to the CLA (middle). Fluorescent CLA terminals are visible in the ACC (right).

### KOR but not MOR and DOR reduce monosynaptic CLA inputs onto L5 ACC PYR cells

Because KOR, MOR, and DOR reduced oEPSCs, we assessed if all three receptors were functionally localized on monosynaptic CLA inputs onto L5 ACC PYR cells. To assess functional opioid receptor expression on CLA terminals projecting onto L5 ACC PYR cells, we pharmacologically isolated monosynaptic transmission with TTX + 4-AP (Fig. 3A). U69 significantly decreased oEPSC amplitude and the antagonist nor-BNI recovered the oEPSC amplitude (% oEPSC reduction; U69: 50.5 ± 10.2%, nor-BNI: 24.9 ± 12.5%, Fig. 3C). DAMGO and DPDPE did not significantly decrease oEPSC amplitude (% oEPSC reduction; DAMGO: 15.6 ± 5.1%, DPDPE: -0.2 ± 5.3%, Fig. 3B, C). These results suggest that KOR but not MOR and DOR are functionally present on CLA inputs onto L5 ACC PYR cells.

**Figure 3.**
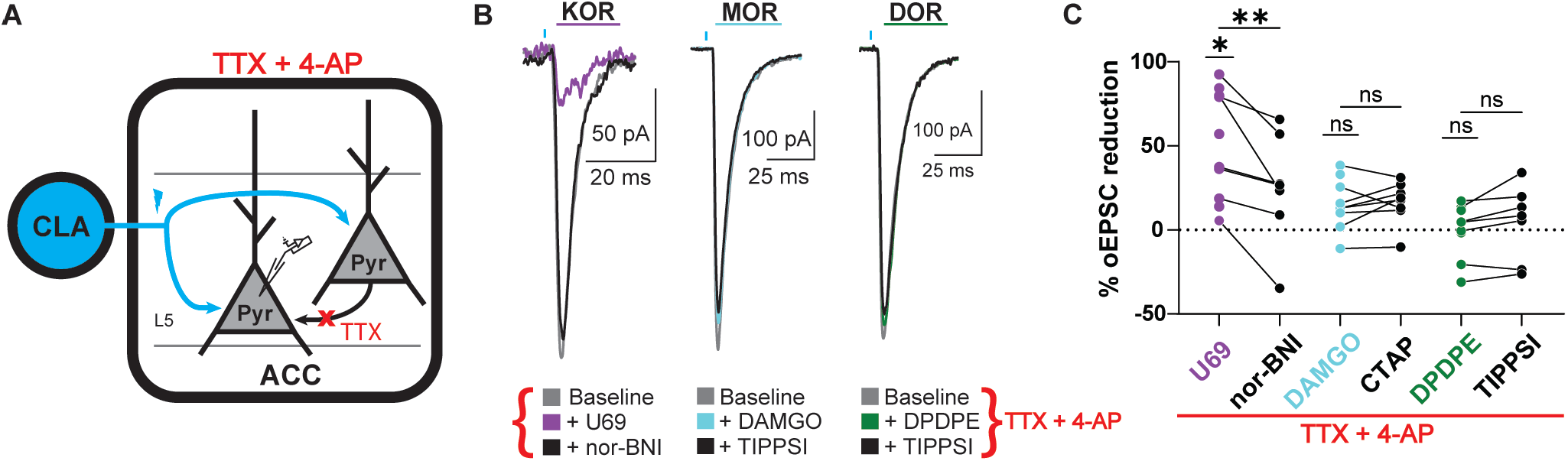
Only KOR activation suppressed monosynaptic oEPSCs from the CLA onto L5 ACC PYR cells. A, Schematic of recording oEPSCs of L5 ACC PYR cells evoked by optical stimulation of CLA input in the presence of TTX + 4-AP. B, Example traces of oEPSCs elicited from optical stimulation of the CLA terminals in the ACC during baseline (gray), left panel: in U69 (1 µM, purple), and in nor-BNI (1 µM, black), middle panel: in DAMGO (1 µM, cyan) followed by CTAP (1 µM, black), and left panel: DPDPE (1 µM, green) followed by TIPPSI (1 µM, black) in the presence of TTX + 4-AP. Blue bars: 1 ms of 470 nm light stimulation. C, Summary data of % oEPSC reduction in agonists and antagonists in (B) relative to baseline oEPSC amplitude: Baseline vs. U69, p = 0.0149, N = 6; U69 vs. norBNI, p = 0.003, N = 6, n = 8; Baseline vs. DAMGO, p = 0.0885, N = 5, n = 9; CTAP: 16.4 ± 4.5 %, DAMGO vs. CTAP, p = 0.6864, N = 5, n = 7; Baseline vs. DPDPE, p = 0.8226, N = 6, n = 9; TIPPSI: 4.5 ± 8.4 %, DPDPE vs. TIPPSI, p = 0.4197, N = 6, n = 6. Mixed-effects analysis with Šídák’s multiple comparisons tests conducted with raw amplitudes. *p ≤ 0.05; **p ≤ 0.01. Data are described as the mean ± standard error of the mean.

### oEPSCs onto L5 ACC PYR cells generated by CLA afferents exhibit both “Early” monosynaptic and “Late” polysynaptic components

Canonical cortical circuits are modeled to amplify incoming input (Douglas and Martin 2004, 2007, Joglekar et al., 2018, Peron et al., 2020). These inputs can generate an early initial excitatory response followed by a late secondary excitatory response mediated through recurrent excitation generated by local excitatory ACC neurons driving the amplification process (Harris and Shepherd 2015, Printz et al., 2023, Vantomme et al., 2025). Therefore, opioid agonists could act at multiple sites to reduce excitatory transmission; directly on CLA terminals in the ACC or on local ACC neurons that are involved in recurrent excitation. Knowing that KOR only reduces monosynaptic CLA inputs, MOR and DOR possibly reduce excitatory polysynaptic transmission driven by the CLA onto L5 ACC PYR cells. To investigate this, we aimed to further characterize the oEPSC by breaking each oEPSC into discrete synaptic events using the 1st temporal derivative of the voltage-clamp waveform. The derivatives of oEPSCs were plotted and an event detection analysis was carried out (Fig. 4A). Baseline events detected from 0 - 10 ms at the time of optical stimulation were binned and plotted. The data were best fit with a double gaussian function (Fig. 4B), indicative of early and late components of the oEPSC. These two components were divided at the trough of the double gaussian curve at 4.72 ms. To determine whether the late component was likely due to polysynaptic recurrent excitation, we isolated the monosynaptic oEPSC, by again recording in TTX and 4-AP (Fig. 4C). oEPSC events derived from baseline oEPSCs averaged 2.8 ± 0.2 events/episode which significantly decreased to 1.2 ± 0.1 events/episode in TTX + 4-AP (Fig. 4D). The majority of these remaining events in TTX + 4-AP fell in the early time window (Fig. 4E, F). As hypothesized, these results suggest that CLA inputs onto L5 ACC PYR cells generate both an early component (< 4.72 ms) in which most monosynaptic oEPSC events fell in and a late component (> 4.72 ms) in which most polysynaptic oEPSC events fell in.

**Figure 4:**
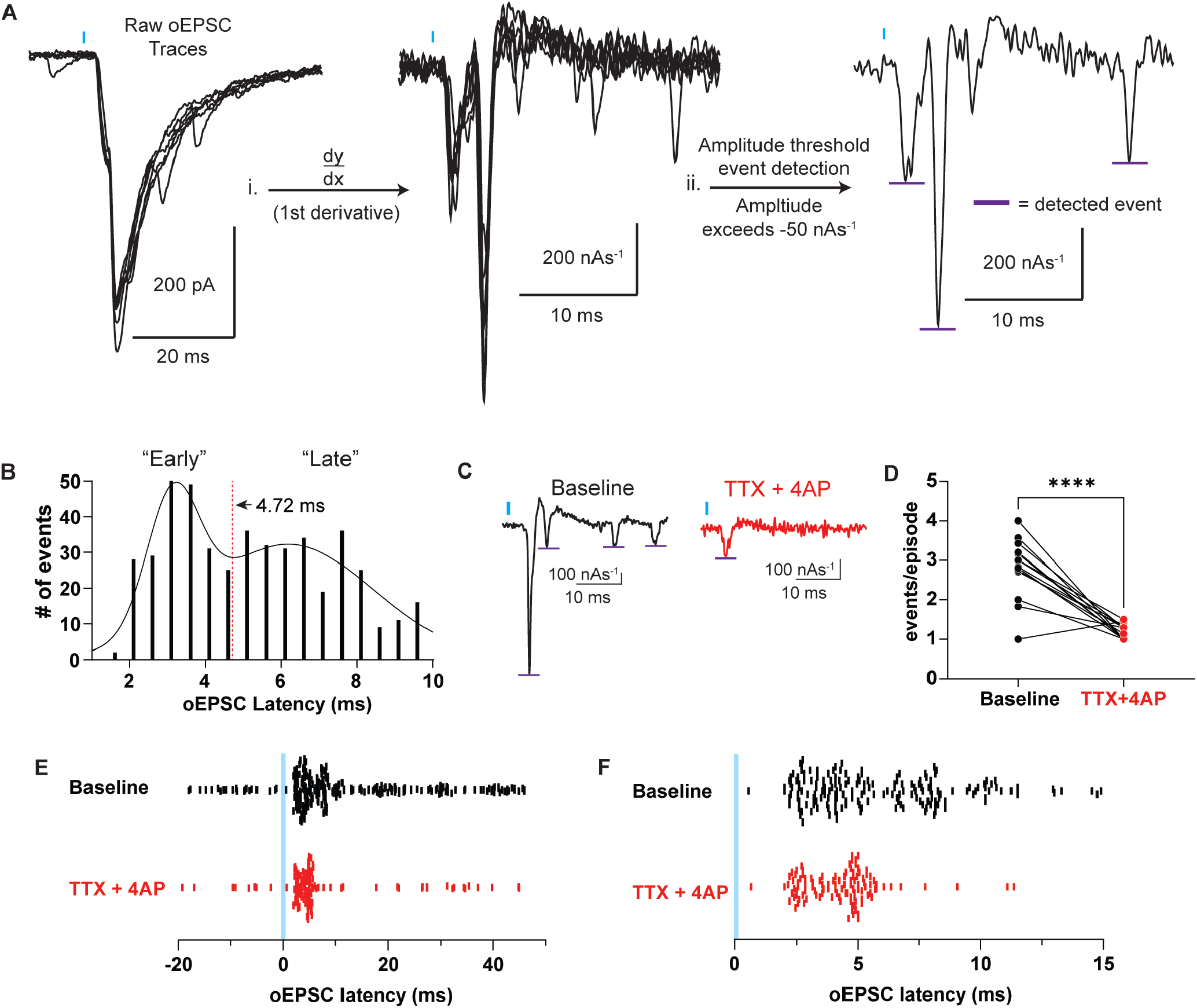
CLA evoked oEPSCs onto ACC L5 pyramidal neurons exhibited an early monosynaptic and late polysynaptic component. A, Workflow: i) differentiating the raw voltage clamp waveform to obtain instantaneous oEPSC slope measurements, ii) event detection analysis with a -50 nA*s-1 event threshold. Purple bars are the maximum slope of the detected events. B, Histogram of time of maximum slope of detected events relative to the optical stimulation (N = 25, n = 52, events detected = 474). C, Example traces of the 1st temporal derivative of voltage clamp oEPSC waveforms during baseline (black) and following TTX + 4-AP (red). Purple lines indicate detected events. D, Comparison of the events/ episode in baseline and TTX + 4-AP conditions (Baseline vs. TTX + 4-AP, p = <0.0001, paired t test). E, Plot of events detected in baseline and TTX + 4-AP conditions from -20 to 50 ms relative to optical stimulation. Each line represents a single event. F, Plot of detected events, as seen in (E), from 0 to 10 ms. Blue bars: 1 ms of 470 nm light stimulation. **** p < 0.0001. Data are represented as the mean ± standard error of the mean.

### MOR agonists suppress the early and late components of the oEPSC while DOR agonists preferentially decrease the late components of the oEPSC

Since MOR and DOR did not reduce monosynaptic excitatory transmission from CLA inputs onto L5 ACC PYR cells, we hypothesized that MOR and DOR might selectively reduce local ACC recurrent excitation. Our data suggest that CLA-ACC oEPSCs are compound oEPSCs composed of both early and late events with late events being predominantly polysynaptic (Fig. 4). To assess the effects of MOR and DOR-mediated reduction of the oEPSC, we differentiated current recordings to obtain the 1st temporal derivative of the voltage clamp waveform (as in Fig. 4) and identified events across baseline and drug conditions. Comparing the cumulative distribution of optically evoked events before and after for either DAMGO (Fig. 5A) or DPDPE (Fig. 5D) revealed a decrease in the number of optically evoked events detected at longer latencies but not shorter latencies. To quantify this observation, the effects of DAMGO and DPDPE on the grouped “early” (< 4.72 ms) and “late” (> 4.72 ms) optically evoked oEPSC events (from Fig. 4B) were analyzed. DAMGO significantly decreased the oEPSC frequency in late oEPSCs but not early oEPSCs (Early oEPSCs events/episode, Baseline: 0.9 ± 0.1, DAMGO: 0.9 ± 0.1; Late oEPSCs events/episode, Baseline: 1.2 ± 0.1, DAMGO: 0.7 ± 0.2, Fig. 5B, C). DPDPE also significantly decreased the oEPSC frequency in late oEPSCs but not early oEPSCs (Early oEPSCs events/episode, Baseline: 0.7 ± 0.2, DPDPE: 0.5 ± 0.1; Late oEPSCs events/episode: Baseline: 1.4 ± 0.2, DPDPE: 1.0 ± 0.2, Fig. 5E, F).

**Figure 5.**
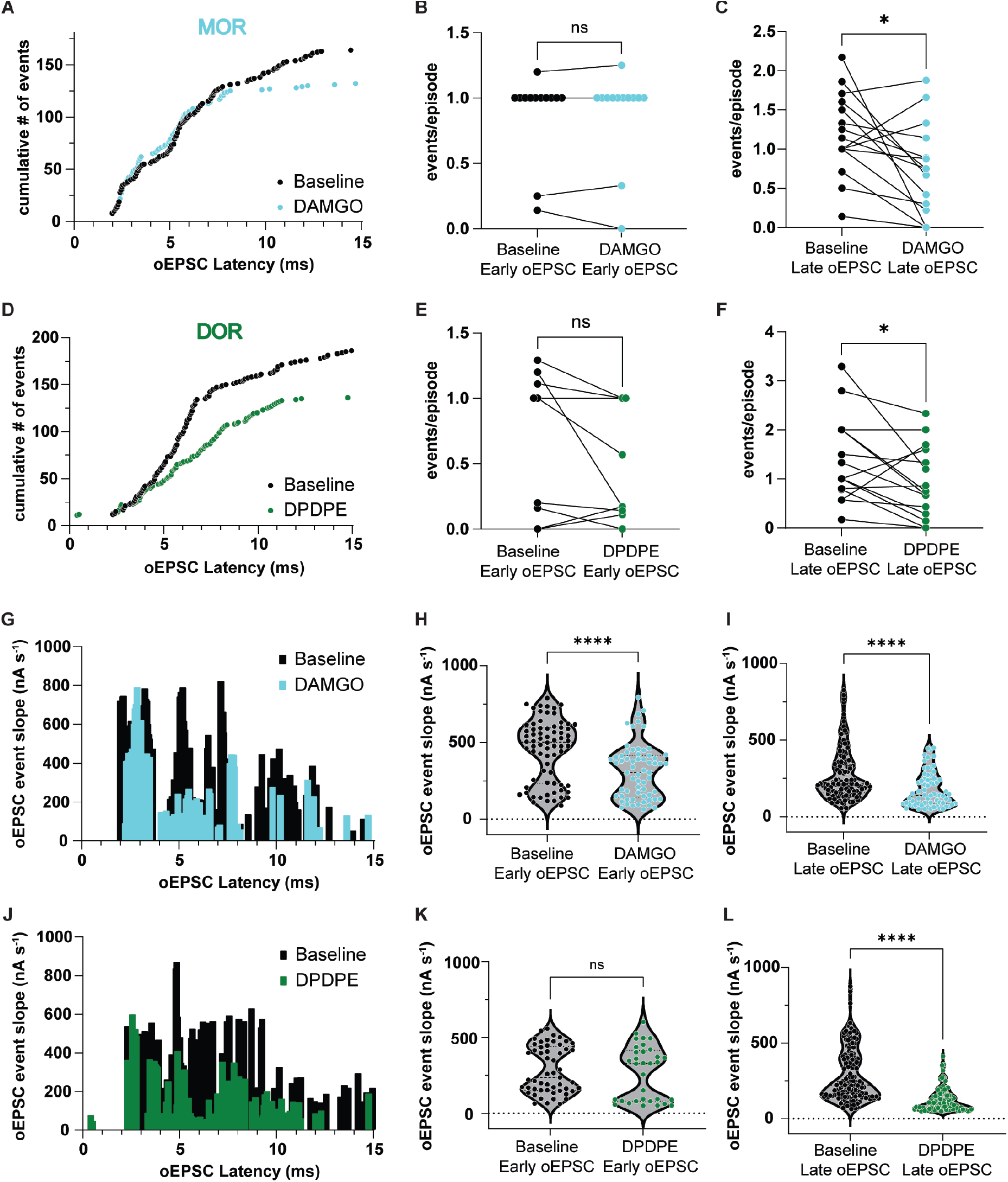
Activation of MOR and DOR alter release frequency and synaptic conductance in a latency and receptor-dependent manner. A, Cumulative distribution of oEPSC events for baseline (black) and DAMGO (cyan) across oEPSC maximum slope latency. B, Summary data of early oEPSC events/episode for baseline and DAMGO conditions (Baseline early oEPSC: 0.9 ± 0.1, DAMGO early oEPSC: 0.9 ± 0.1, Baseline vs. DAMGO, p = 0.9555, paired t-test, N = 16, n = 14). C, Summary data of late oEPSC events/episode for Baseline and DAMGO conditions (Baseline late EPSC: 1.2 ± 0.1, DAMGO late oEPSC: 0.7 ± 0.2, Baseline vs. DAMGO, p = 0.0118, paired t-test, N = 16, n = 16). D, Cumulative distribution of oEPSC events for baseline (black) and DPDPE (green) across oEPSC maximum slope latency. E, Summary data of events/episode of early oEPSCs for baseline and DPDPE conditions (Baseline early oEPSC: 0.7 ± 0.2, DPDPE early oEPSC: 0.5 ± 0.1, Baseline vs. DPDPE, p = 0.1361, paired t-test, N = 13, n = 10). F, Summary data of late oEPSC events/episode for baseline and DPDPE conditions (Baseline late oEPSC: 1.4 ± 0.2, DAMGO late oEPSC: 0.9 ± 0.2, Baseline vs. DPDPE, p = 0.0427, paired t-test, N = 13, n = 15). G, Histogram of oEPSC event slope vs. oEPSC **latency for events detected in baseline** (black) and DAMGO (cyan) conditions. H, Violin plot of maximum oEPSC slope of pooled early oEPSC events in baseline and DAMGO conditions (Baseline early oEPSC: 451.3 ± 23.7 nA*s-1, DAMGO early oEPSC: 295.4 ± 25.5 nA*s-1, Baseline vs. DAMGO, p = <0.0001, unpaired t-test, N = 16, n = 18). I, Violin plot of maximum oEPSC slope of pooled late oEPSC events in baseline and DAMGO conditions (Baseline late oEPSC: 273.0 ± 15.5 nA*s-1, DAMGO late oEPSC: 175.8 ± 13.7 nA*s-1, Baseline vs. DAMGO, p = <0.0001, unpaired t-test, N = 16, n = 18). J, Histogram oEPSC event slope vs. oEPSC latency for events detected in baseline (black) and DPDPE (green) conditions. K, Violin plot of maximum oEPSC slope of pooled early oEPSC events in baseline and DPDPE conditions (Baseline early oEPSC: 286.8 ± 21.2 nA*s-1, DAMGO early oEPSC: 268.3 ± 31.0 nA*s-1, Baseline vs. DAMGO, p = 0.6131, unpaired t-test, N = 13, n = 17). L, Violin plot of maximum oEPSC slope of pooled late oEPSC events in baseline and DPDPE conditions (Baseline late oEPSC: 287.6 ± 14.2 nA*s-1, DPDPE late oEPSC: 124.0 ± 7.9 nA*s-1, Baseline vs. DPDPE, p = <0.0001, unpaired t-test, N = 13, n = 17). *p ≤ 0.05; **** p < 0.0001. Data are represented as the mean ± standard error of the mean.

Next, we compared the maximum slope of each oEPSC event across conditions, which is related to synaptic conductance (Johnston and Wu 1994). In this instance, DAMGO significantly decreased the average oEPSC event slope in both early oEPSCs and late oEPSCs (Early oEPSC event slope, Baseline: 451.3 ± 23.7 nA*s-1, DAMGO: 295.4 ± 25.5 nA*s-1; Late oEPSC event slope, Baseline: 273.0 ± 15.5 nA*s-1, DAMGO: 175.8 ± 13.7 nA*s-1; Fig. 5H, I). However, DPDPE only significantly decreased the late oEPSCs but not the early oEPSCs (Early oEPSC event slope, Baseline: 286.8 ± 21.2 nA*s-1, DPDPE: 268.3 ± 31.0 nA*s-1; Late oEPSC event slope, Baseline: 287.6 ± 14.2 nA*s-1, DPDPE: 124.0 ± 7.9 nA*s-1; Fig. 5K, L). Holding current did not change in the presence of the agonist or antagonist. These data suggest that MOR and DOR reduced the late polysynaptic component and early monosynaptic components of the oEPSC differently, with MOR affecting both, while DOR primarily inhibited the late polysynaptic oEPSC. Further, MOR affected early and late events differently; early events had a shallower maximum slope, while late events had both decreased frequency and shallower maximum slope. These results support the idea that MOR and DOR primarily suppress polysynaptic transmission in the CLA-ACC pathway; however, MOR reduced the postsynaptic conductance of early oEPSCs, suggesting a particular role of modulating but not completely inhibiting monosynaptic inputs; however, Fig. 3C suggested little to no MOR functional expression on monosynaptic CLA inputs onto L5 ACC PYR cells.

### MOR and DOR reduce excitation in a latency dependent manner whereas KOR reduces all excitation regardless of latency

Our data suggested that KOR and possibly MOR reduced early monosynaptic CLA inputs onto ACC L5 PYR cells whereas both MOR and DOR reduced late recurrent excitation generated by CLA inputs. We showed that late oEPSCs that are blocked by TTX are primarily polysynaptic (Fig. 4); however, we wanted to observe if this effect was still present with both monosynaptic and polysynaptic excitatory transmission combined in our analysis. Opioid agonists acting directly on CLA terminals should suppress both the early initial excitatory response and the late slower recurrent excitation due to reduced excitatory drive into the ACC. Opioid agonists acting on local ACC neurons should preferentially reduce late recurrent excitation while sparing the early excitatory responses. To investigate this, we used the raw oEPSC data seen in (Fig. 2F). First, we measured the oEPSC onset latency from raw traces under baseline conditions for all recordings. Similar to the findings from the derivative analysis in Fig. 4B, the oEPSC latency histogram revealed a bimodal distribution of oEPSC onset latency that was best fit with a double gaussian function, suggesting two different oEPSC populations identified based on onset latency (Fig. 6A, B). The trough of the double gaussian fit occurred at 4.25 ms; therefore, oEPSCs were grouped into “early” oEPSC’s (onset values less than 4.25 ms) and “late” oEPSC’s (onset values greater than 4.25 ms) (Fig. 6B, C). If KOR acts on monosynaptic CLA inputs and MOR and DOR act on polysynaptic inputs, then there should only be a positive correlation between oEPSC onset latency and MOR and DOR reduction of the oEPSC, respectively. Furthermore, since KOR acts on monosynaptic CLA inputs, both early and late oEPSCs should be suppressed due to inhibiting CLA terminals which are then unable to activate recurrent excitation.

**Figure 6.**
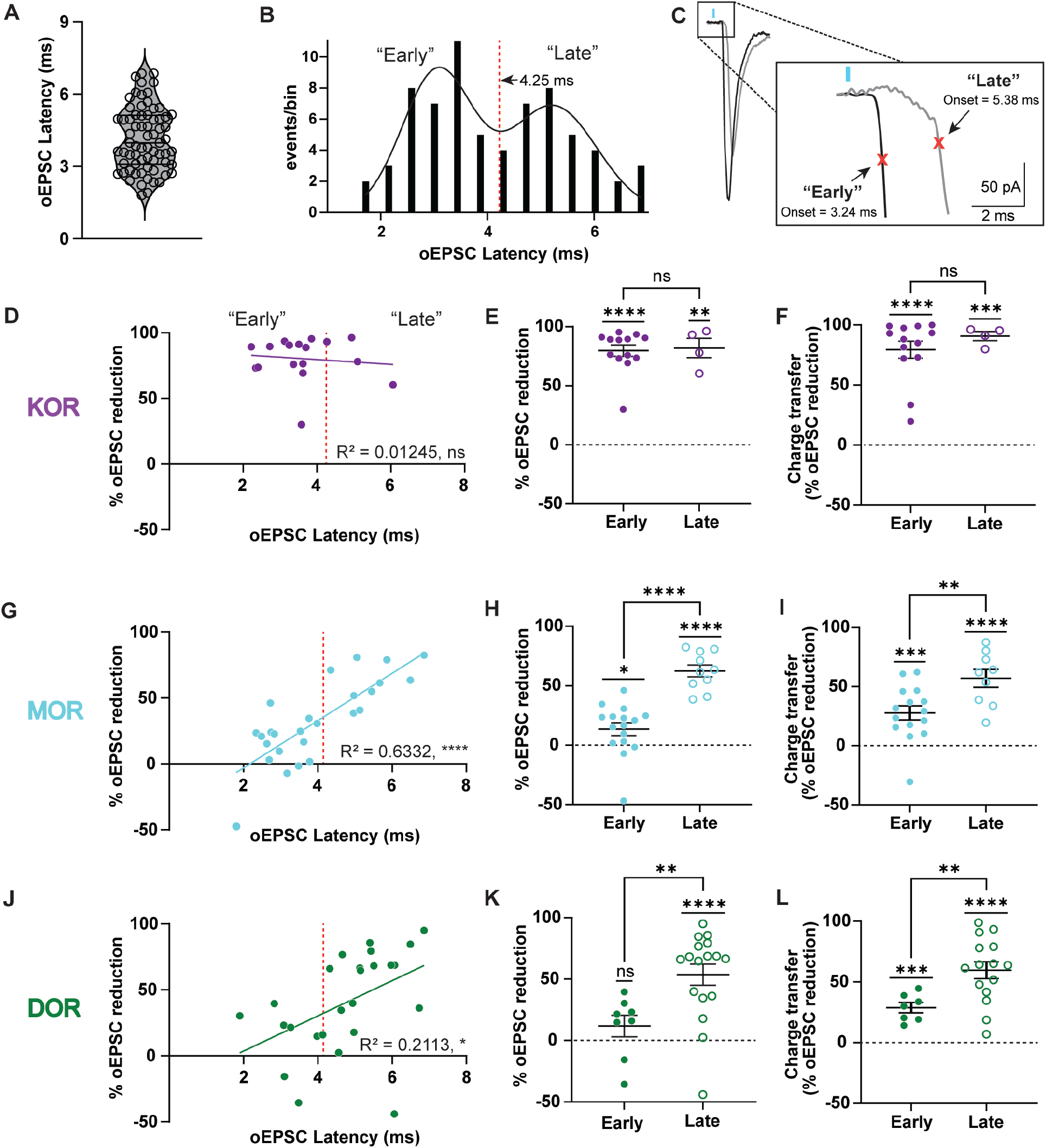
oEPSC onset latency determines oEPSC inhibition by MOR and DOR but not KORagonists. A, Violin plot of individual baseline oEPSC latencies (n = 69). B, Histogram of values as shown in (A). C, Example traces of a early (blue) and late oEPSC (grey). Traces are from two different cells. D, Plot of the % oEPSC reduction by U69 relative to oEPSC latency. There was no correlation between the % oEPSC reduction by U69 and oEPSC onset latency (Simple linear regression, R2 = 0.01245, p = 0.6698). E, Summary data of % oEPSC reduction by U69 in early and late oEPSC populations shown in (D) (early oEPSCs: 79.8 ± 4.8 %, early oEPSCs vs. 0, p = <0.0001, n = 13; late oEPSCs: 82.0 ± 8.2 %, late oEPSCs vs. 0, p = 0.0021, n = 4, one sample t-test, early vs. late, p = 0.8244, unpaired t-test). F, Summary data of % oEPSC reduction of charge transfer by U69 in early and late oEPSC populations (early oEPSCs: 79.5 ± 7.0 %, early vs. 0, p = <0.0001; late oEPSCs: 90.8 ± 3.8 %, late oEPSCs vs. 0, p = 0.0002, one sample t-test, early vs. late, p = 0.4030, unpaired t-test). G, Plot of the MOR effects on % oEPSC reduction by DAMGO relative to oEPSC latency. There was a significant correlation between the % oEPSC reduction by DAMGO and oEPSC onset latency (Simple linear regression, R2 = 0.6332, p = <0.0001). H, Summary data of % oEPSC reduction by DAMGO in early and late oEPSC populations (G) (early oEPSCs: 13.5 ± 5.5 %, early oEPSCs vs. 0, p = 0.0257, n = 16; late oEPSCs: 62.4 ± 5.0 %, late oEPSCs vs. 0, p = <0.0001, n = 10, one sample t-test, early vs. late, p = <0.0001, unpaired t-test). I, Summary data of % oEPSC reduction of charge transfer by DAMGO in early and late oEPSC populations (early oEPSCs: 27.8 ± 6.0 %, early oEPSCs vs. 0, p = 0.0004, late oEPSCs: 57.0 ± 7.5 %, late oEPSCs vs. 0, p = <0.0001, one sample t-test, early vs. late, p = 0.0065, unpaired t-test). J, Plot of the DOR effects on % oEPSC reduction by DPDPE relative to oEPSC latency. There was a significant correlation between the % oEPSC reduction by DPDPE and oEPSC onset latency (Simple linear regression, R2 = 0.2113, p = 0.0208). K, Summary data of % oEPSC reduction by DPDPE in early and late oEPSC populations (J) (early oEPSCs: 11.8 ± 8.8 %, early oEPSCs vs. 0, p = 0.2242, n = 8, late oEPSCs: 53.6 ± 8.6 %, late oEPSCs vs. 0, p = <0.0001, n = 17, one sample t-test, early vs. late, p = 0.0067, unpaired t-test). L, Summary data of % oEPSC reduction of charge transfer by DPDPE in early and late oEPSC populations (early oEPSCs: 28.8 ± 4.3 %, early oEPSCs vs. 0, p = 0.0005, late oEPSCs: 59.7 ± 6.9 %, late oEPSCs, vs. 0, p = <0.0001, one sample t-test, early vs. late, p = 0.0085, unpaired t-test. *p ≤ 0.05; ** p ≤ 0.01; *** p ≤ 0.001; **** p < 0.0001. Dashed red line: 4.25 ms. Data are represented as the mean ± standard error of the mean.

As hypothesized, there was no significant correlation between oEPSC onset latency and oEPSC inhibition by U69 (Fig. 6D). U69 significantly reduced the oEPSC peak amplitude of both early and late onset oEPSCs (Early % oEPSC reduction: 79.8 ± 4.8%, Late % oEPSC reduction: 82.0 ± 8.2%, Fig. 6E). In contrast, there was a significant linear correlation between oEPSC onset latency and % oEPSC reduction by DAMGO and DPDPE (DAMGO: R2 = 0.6, Fig. 6G; DPDPE: R2 = 0.2, Fig. 6J). While DAMGO inhibited both early and late onset oEPSCs, the late oEPSCs were reduced significantly more than the early onset oEPSCs (Early % oEPSC reduction: 13.5 ± 5.5%, Late % oEPSC reduction: 62.4 ± 5.1%, Fig. 6H). DPDPE application did not significantly inhibit early onset oEPSCs but did inhibit late onset EPSCs (Early % oEPSC reduction: 11.8 ± 8.8%, Late % oEPSC reduction: 53.6 ± 8.6%, Fig. 6K).

Because all three agonists decreased the late individual oEPSC events and late onset oEPSCs, they should all decrease the total charge transfer resulting from compound oEPSCs. As expected, U69, DAMGO, and DPDPE all significantly inhibited the charge transfer from both early and late onset oEPSCs. U69 reduced the charge transfer of both early and late oEPSCs efficaciously and to a similar degree (Early % oEPSC reduction: 79.5 ± 7.0%, Late % oEPSC reduced: 90.8 ± 3.8%, Fig. 6F). DAMGO and DPDPE reduced the charge transfer of both early and late onset oEPSCs but the late onset oEPSCs were significantly more reduced (DAMGO; Early % oEPSC reduction: 27.8 ± 6.0%, Late % oEPSC reduction: 57.0 ± 7.5%, Fig. 6I, DPDPE: Early % oEPSC reduction: 28.8 ± 4.3%, Late % oEPSC reduction: 59.7 ± 6.9%, Fig. 6L). Together, these results suggest that opioid receptor modulation acts on different pathways along the CLA-ACC circuit, where MOR and DOR activation preferentially suppresses late oEPSCs while KOR activation drastically suppresses all oEPSCs. These results are consistent with KOR being expressed on CLA terminals in the ACC while MOR and DOR are expressed on local ACC neurons that modulate recurrent excitation. Consistent with Fig. 5, specific MOR-mediated reduction of the early component of the oEPSC is also seen in Fig. 6H, I, supporting the idea that MOR could partially or indirectly suppress monosynaptic CLA inputs.

## Discussion

This study characterized functional opioid receptor expression of the CLA-ACC circuit. While multiple studies have explored the relevance of this circuit in pain processing, they have not explicitly addressed the involvement of opioids and their modulation of neurotransmission. Here, we used spatial transcriptomics, slice electrophysiology, optogenetics, and pharmacology to determine where and how opioid receptor activation affects excitatory synaptic transmission from the CLA to ACC L5 PYR cells. We found that excitatory transmission from the CLA was inhibited by KOR, MOR, and DOR agonists. Activation of KOR strongly suppressed excitatory monosynaptic inputs from the CLA, whereas MOR and DOR had very little or no effect on these monosynaptic inputs. This suggests that KOR is expressed on CLA terminals within the ACC and activation of KOR can virtually eliminate transmission from the CLA to the ACC (Fig. 7). On the contrary, MOR and DOR agonists appear to preferentially suppress polysynaptic excitation from the CLA to ACC L5 PYR cells (Fig. 7); late onset latency oEPSCs were suppressed by these agonists while early onset latency oEPSCs were mostly spared. Therefore, inhibition of specific aspects of the CLA-ACC circuit by all three opioid receptor types may serve to shape the temporal dynamics of network-level processing within the ACC. Our results suggest that opioid effects on circuits involved in pain and consciousness can be shaped by a diverse activation of opioid receptors (Navratilova et al., 2024).

**Figure 7:**
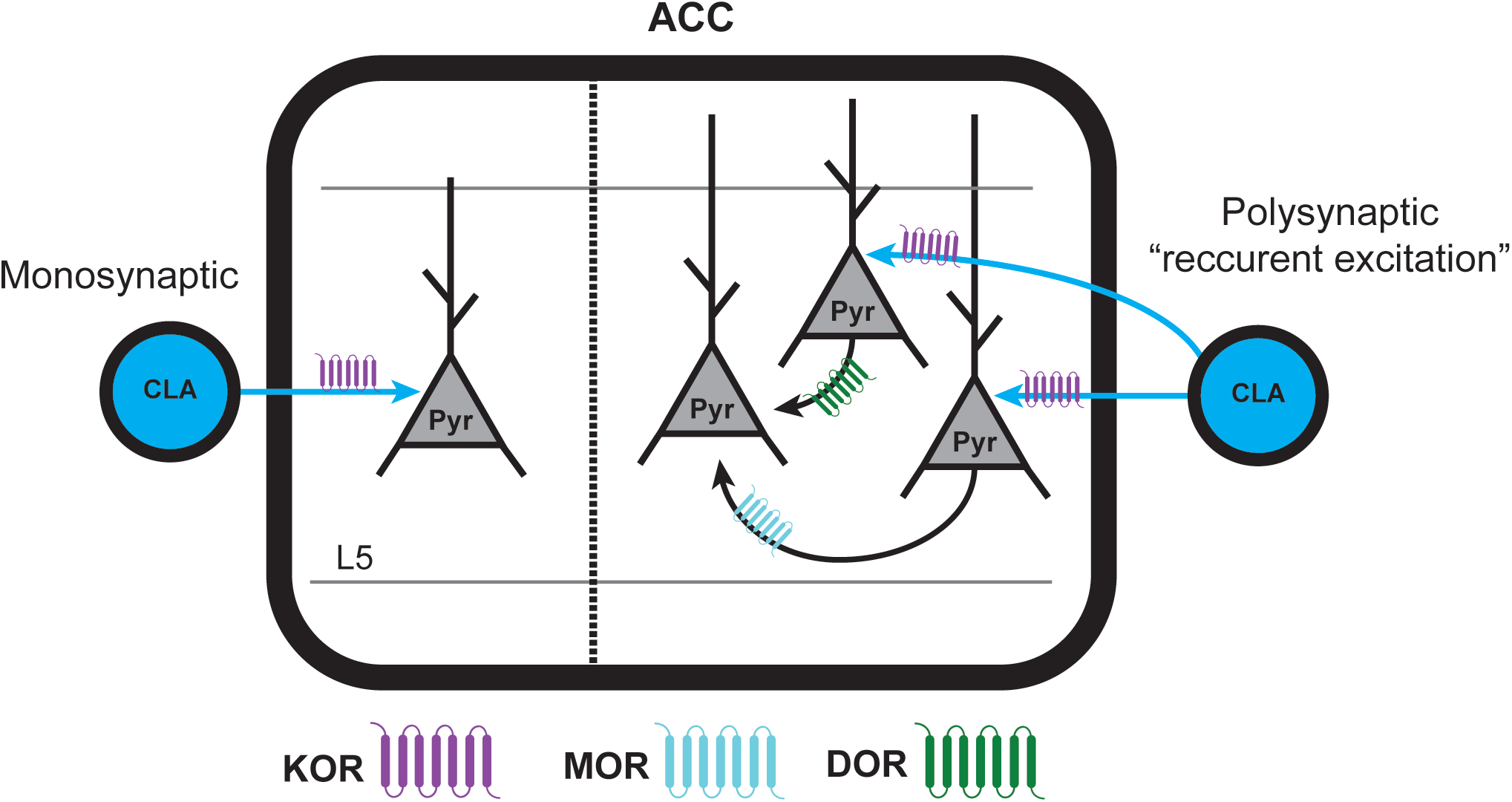
Proposed model of how excitatory transmission is modulated by opioid receptors along the CLA-ACC pathway. Left: Schematic of KOR (purple) reducing excitatory monosynaptic inputs from the CLA onto L5 ACC PYR cells. Right: Schematic of MOR (cyan) and DOR (green) reducing recurrent excitation in the ACC generated by CLA inputs.

While it has been shown that KOR, MOR, and DOR transcripts are expressed along the CLA-ACC pathway, functional expression and their role on modulating synaptic transmission was previously unknown (Tempel and Zukin 1987, George et al., 1994, Mansour et al., 1994, Georges et al., 1998, Wang et al., 2023). We found that all three opioid receptors modulate excitatory synaptic transmission onto L5 ACC PYR cells. Surprisingly, we found that MOR and DOR reduce the polysynaptic component of the oEPSC whereas KOR reduces the monosynaptic component. This aligns with previously understood expression patterns since KOR is highly expressed and MOR is moderately expressed in the CLA whereas DOR exhibits high expression and MOR exhibits moderate expression in the ACC. We have multiple lines of evidence that suggest KOR is functionally expressed on CLA afferents and that KOR activation dramatically reduces glutamate release from these terminals. While we did not study KOR expression on ACC neurons, our data is consistent with high levels of KOR expression on CLA inputs onto excitatory ACC neurons. However, we can’t rule out KOR expression on neurons within the ACC.

The mRNA transcript for MOR, Oprm1, is highly expressed in ACC pyramidal cells with highest expression in L6 (James et al., bioRxiv). Recurrent excitation driven by pyramidal-pyramidal cell interactions can be inhibited if MOR is functionally expressed on the synaptic terminals. This could explain the latency-dependent reduction of oEPSCs by MOR. Future studies identifying functional MOR expression in pyramidal cells in the ACC is needed to evaluate MOR-mediated reduction of local recurrent excitation.

Multiplexed error-robust fluorescence in situ hybridization (MERFISH) data from the Allen Institute for Brain Science shows robust expression of DOR in L2/3 and L6 ACC PYR cells (Yao et al., 2023). In rats, DOR activation has been shown to hyperpolarize some ACC pyramidal cells and inhibit EPSCs onto ACC pyramidal cells (Tanaka and North 1994). If DOR is functionally expressed in ACC PYR cells, then DOR activation would reduce recurrent excitation in the ACC. This hypothesis would be supported by our data showing that DOR suppresses the late oEPSC but not the early oEPSC. Further, DOR does reduce the early oEPSC component of compound oEPSCs suggesting that DOR suppresses later polysynaptic oEPSC events.

In agreement with our interpretation that KOR activation reduces monosynaptic input whereas MOR and DOR activation reduces polysynaptic input from the CLA, we observed that only U69 suppressed the oEPSC in TTX + 4-AP conditions. However, MOR activation did slightly reduce early oEPSCs (Fig. 6 G-I) and decreased the early oEPSC event slope (Fig. 5H) but not the frequency of fast oEPSC events (Fig. 4B), suggesting MORs may affect monosynaptic CLA oEPSCs (directly or indirectly). Additionally, in the presence of TTX + 4-AP, MOR activation showed a trend towards slightly inhibiting the oEPSC, however this did not reach statistical significance. 4-AP (in the absence of TTX) has been shown to blunt the ability of MOR agonists to reduce GABA transmission in some contexts (Vaughn et al., 1997, Ingram et al., 1998), making a lack of reduction in the presence of 4-AP somewhat difficult to interpret. Nonetheless, even at saturating concentrations of the full agonist DAMGO, there was, at most, only modest inhibition of the early monosynaptic oEPSCs when measured using multiple approaches. This is in spite of the moderate uniform level of oprm1 expression across all the glutamatergic CLA neurons (Fig. 1B, C). This suggests that the moderate levels of oprm1 transcript do not result in functional MOR expression or that MOR expressed in CLA neurons is not transported to CLA terminals in the ACC in high abundance. Altogether, these data suggest that Kappa opioid receptors may serve as a gate keeper for information flow from the CLA into the ACC while Mu, and Delta opioid receptors shape the temporal dynamics of ACC processing.

By activating different complements of opioid receptors, endogenous opioids have the potential to shape cortical processing. Dynorphin, the KOR-preferring opioid peptide, can modulate monosynaptic transmission of CLA inputs into the ACC resulting in filtering or gating specific sensory inputs and potentially having a role in pain processing (Wang et al., 2023). Local ACC neurons, such as somatostatin inhibitory interneurons, express dynorphin (Tremblay et al., 2016) and can act on these KOR-sensitive inputs, shunting incoming information. Enkephalin, the DOR/MOR-preferring peptide, may modulate polysynaptic transmission including slower, sustained excitatory feedback loops in the ACC, potentially modulating more cognitive or attentional aspects of sensory information. Beta-endorphin, the MOR-preferring peptide, may preferentially reduce longer latency oEPSCs, while sparing early excitatory oEPSCs. Together, dynorphins, enkephalins and endorphins likely collaborate to finely tune network-level dynamics in the CLA-ACC circuit, balancing immediate sensory input processing with sustained, complex cognitive or affective processing, which could have implications for understanding pain and consciousness.

The CLA-ACC circuit has been demonstrated to play roles in pain (Xu et al., 2022, Ntamati et al., 2023, Koga et al., 2024, Zhang and Zamponi 2024), attention (Mathur 2014, Goll et al., 2015), and saliency (Remedios et al., 2010, Kitanishi and Matsuo 2017, Reser et al., 2017). Therefore, some aspects of analgesic, attentive, and salient behaviors are likely to be modulated by opioids. Given the strong opioid receptor modulation within CLA-ACC circuitry, future research should seek to understand how these different receptors alter information processing in these different brain states and behavioral contexts.

## Acknowledgements

This work was supported by National Institutes on Drug Abuse Grants R01DAG042779 (to W.T.B.), and T32DA007281 (to J.M.R.). We thank Dr. Pierre Apostolides for guidance and comments on the manuscript. We thank Dr. Erica Levitt and Dr. Paul Kramer for comments on this manuscript.

